# The M2 Gene Is a Determinant of Reovirus-Induced Myocarditis

**DOI:** 10.1101/2021.07.16.452757

**Authors:** Marcelle Dina Zita, Matthew B. Phillips, Johnasha D. Stuart, Asangi R. Kumarapeli, Anthony J. Snyder, Amairani Paredes, Vijayalakshmi Sridharan, Marjan Boerma, Pranav Danthi, Karl W. Boehme

**Affiliations:** Department of Microbiology and Immunology, University of Arkansas for Medical Sciences; Department of Pathology, University of Arkansas for Medical Sciences; Department of Biology, Indiana University-Bloomington, University of Arkansas for Medical Sciences; Department of Pharmaceutical Sciences, University of Arkansas for Medical Sciences; Winthrop P. Rockefeller Cancer Institute, University of Arkansas for Medical Sciences; Center for Microbial Pathogenesis and Host Inflammatory Responses, University of Arkansas for Medical Sciences

## Abstract

Although a broad range of viruses cause myocarditis, the mechanisms that underlie viral myocarditis are poorly understood. Here, we report that the M2 gene is a determinant of reovirus myocarditis. The M2 gene encodes outer capsid protein μ1, which mediates host membrane penetration during reovirus entry. We infected newborn C57BL/6 mice with reovirus strain type 1 Lang (T1L) or a reassortant reovirus in which the M2 gene from strain type 3 Dearing (T3D) was substituted into the T1L genetic background (T1L/T3DM2). T1L was non-lethal in wild-type mice, whereas greater than 90% of mice succumbed to T1L/T3DM2 infection. T1L/T3DM2 produced higher viral loads than T1L at the site of inoculation. In secondary organs, T1L/T3DM2 was detected with more rapid kinetics and reached higher peak titers than T1L. We found that hearts from T1L/T3DM2-infected mice were grossly abnormal, with large lesions indicative of substantial inflammatory infiltrate. Lesions in T1L/T3DM2-infected mice contained necrotic cardiomyocytes with pyknotic debris, and extensive lymphocyte and histiocyte infiltration. In contrast, T1L induced the formation of small purulent lesions in a small subset of animals, consistent with T1L being mildly myocarditic. Finally, more activated caspase-3-positive cells were observed in hearts from animals infected with T1L/T3DM2 compared to T1L. Together, our findings indicate that substitution of the T3D M2 allele into an otherwise T1L genetic background is sufficient to change a non-lethal infection into a lethal infection. Our results further indicate that T3D M2 enhances T1L replication and dissemination *in vivo*, which potentiates the capacity of reovirus to cause myocarditis.

**IMPORTANCE:** Reovirus is a non-enveloped virus with a segmented double-stranded RNA genome that serves as a model for studying viral myocarditis. The mechanisms by which reovirus drives myocarditis development are not fully elucidated. We found that substituting the M2 gene from strain type 3 Dearing (T3D) into an otherwise type 1 Lang (T1L) genetic background (T1L/T3DM2) was sufficient to convert the non-lethal T1L strain into a lethal infection in neonatal C57BL/6 mice. T1L/T3DM2 disseminated more efficiently and reached higher maximum titers than T1L in all organs tested, including the heart. T1L is mildly myocarditic and induced small areas of cardiac inflammation in a subset of mice. In contrast, hearts from mice infected with T1L/T3DM2 contained extensive cardiac inflammatory infiltration and more activated caspase-3-positive cells, which is indicative of apoptosis. Together, our findings identify the reovirus M2 gene as a new determinant of reovirus-induced myocarditis.

## INTRODUCTION

Numerous viruses, such as cytomegalovirus (CMV), coxsackievirus B3 (CVB3), parvovirus B19, and severe acute respiratory syndrome coronavirus-2 (SARS-CoV-2) can cause myocarditis, which is characterized by inflammation of the heart muscle that can lead to cardiac damage, electrophysiological abnormalities, heart failure, or sudden death (1). Viral myocarditis is driven by a combination of virus-induced cytopathic effects in cardiac cells and the host inflammatory response to infection (2–5). Although a significant public health risk, many open questions remain with respect to the viral factors and host mechanisms that underlie viral myocarditis.

Mammalian orthoreovirus (reovirus) is a non-enveloped, double-stranded RNA (dsRNA) virus with a segmented genome that serves as a model for studying viral myocarditis (6). Some, but not all, reovirus strains cause myocarditis in neonatal mice (7, 8). The segmented nature of the reovirus genome has enabled the identification of specific genes that associate with myocarditis induction. Such studies have identified proteins that comprise the viral inner shell (core) as determinants of myocarditis (7, 9, 10). Yet, amino acid changes in core proteins of myocarditic and non-myocarditic strains that define differences in heart disease have remain unidentified. Interestingly, most myocarditic strains are reassortants. Clone 8B, a highly myocarditic reovirus strain, was generated via reassortment between the mildly myocarditic reovirus strain type 1 Lang (T1L) and the non-myocarditic strain type 3 Dearing (T3D) (11). Clone 8B contains 8 gene segments from T1L in combination with the S1 and M2 genes from T3D (11). The reovirus S1 gene encodes viral attachment protein σ1 and non-structural protein σ1s, while M2 encodes outer capsid protein μ1 (6). T1L/T3Dμ1σ3 contains 8 gene segments from T1L with the M2 and S4 genes from T3D (12). The S4 gene encodes outer capsid protein σ3 that complexes with μ1 on the reovirus virion (6). The KC19 strain that contains 7 genes from T1L and the S1, S3 (encoding nonstructural protein σNS), and M2 genes from T3D also has higher myocarditic potential than T1L (7). Common among clone 8B, T1L/T3Dμ1σ3, and KC19 is that they contain the M2 gene from T3D. This observation raises the possibility that the T3D-derived M2 gene impacts the myocarditic potential of reovirus strains. Yet, this idea has not been formally tested.

Reovirus infection is initiated by binding of viral attachment protein σ1 to cell surface receptors, followed by internalization of the particle by receptor-mediated endocytosis (13, 14). In the endosome, heterohexameric complexes of μ1 and σ3 that form the viral outer capsid are cleaved by the activity of endosome-resident cathepsin proteases to facilitate virus entry (15–17). The σ3 protein is degraded, and μ1 is cleaved to enable conformational changes in μ1 required for endosomal membrane penetration and release viral cores into the cytoplasm for transcription and subsequent replication (6, 15–17). In addition to its role in entry, the μ1 protein also contributes to induction of apoptosis in reovirus infected cells (6, 18, 19). T3 and T1 reoviruses differ in many aspects of virus replication and host response. Among them, several phenotypes are associated with T3-derived M2. A single-gene reassortant containing the M2 gene from T3D strain in an otherwise T1L genetic background (T1L/T3DM2) has increased attachment to host cells compared to T1L (20). Enhanced binding of T1L/T3DM2 correlated with greater infectivity and cell death induction in cultured cells. These observations suggest that the reovirus M2 gene, and by extension the μ1 protein, can influence reovirus tropism and disease *in vivo*. The presence of T3D M2 also allows more efficient conformational changes that occur during cell entry. Consequently, membrane-penetration and core delivery is also more efficient for T3D M2 containing viruses (21–23). The higher apoptotic potential of T3 strains in comparison to T1 strains also maps to the μ1-encoding M2 gene segment (18, 19). The precise function of μ1 in apoptosis induction is unclear since seemingly contradictory data exist to support the role both incoming μ1 from capsids and newly synthesized μ1 produced during virus replication (18, 24–26). Nonetheless, mutations in the □ domain of T3D μ1 diminish the apoptotic capacity of reovirus *in vivo* which is associated with attenuated virulence (27).

Here, we assessed the function of M2 in reovirus pathogenesis. We found that in contrast to T1L, T1L/T3DM2 was lethal in neonatal mice. T1L/T3DM2 replicated to higher titers than T1L in all organs tested. Hearts from mice infected with T1L/T3DM2 displayed signs of overt myocarditis, including extensive purulent lesions with marked inflammatory infiltrate. In contrast, hearts from T1L-infected animals appeared largely normal or had small lesions which is consistent with T1L being mildly myocarditic. Together, these findings indicate that the T3D M2 gene is a determinant of reovirus-induced myocarditis.

## RESULTS

### T1L/T3DM2 is more virulent than T1L in neonatal mice

To determine whether the T3D M2 allele modulates reovirus virulence, we assessed survival of neonatal C57BL/6 mice infected orally with T1L or T1L/T3DM2 viruses (Fig 1A). Consistent with previous findings (6), all mice inoculated with T1L survived. In contrast, 90% of mice succumbed to infection with T1L/T3DM2. Similar results were obtained with intracranial inoculation (Fig 1B). Following T1L infection, 11 of 12 mice survived (91.7%), whereas 1 of 14 mice survived (7.1%) T1L/T3DM2 infection. These data indicate that replacing with T1L M2 gene with the T3D M2 allele converts the non-lethal T1L strain into a lethal infection in neonatal mice. Further, because infection with T1L/T3DM2 was lethal following both peroral and intracranial inoculation, our work indicates that the effect of T3D M2 on reovirus virulence is independent of the inoculation route.

**Figure 1.**
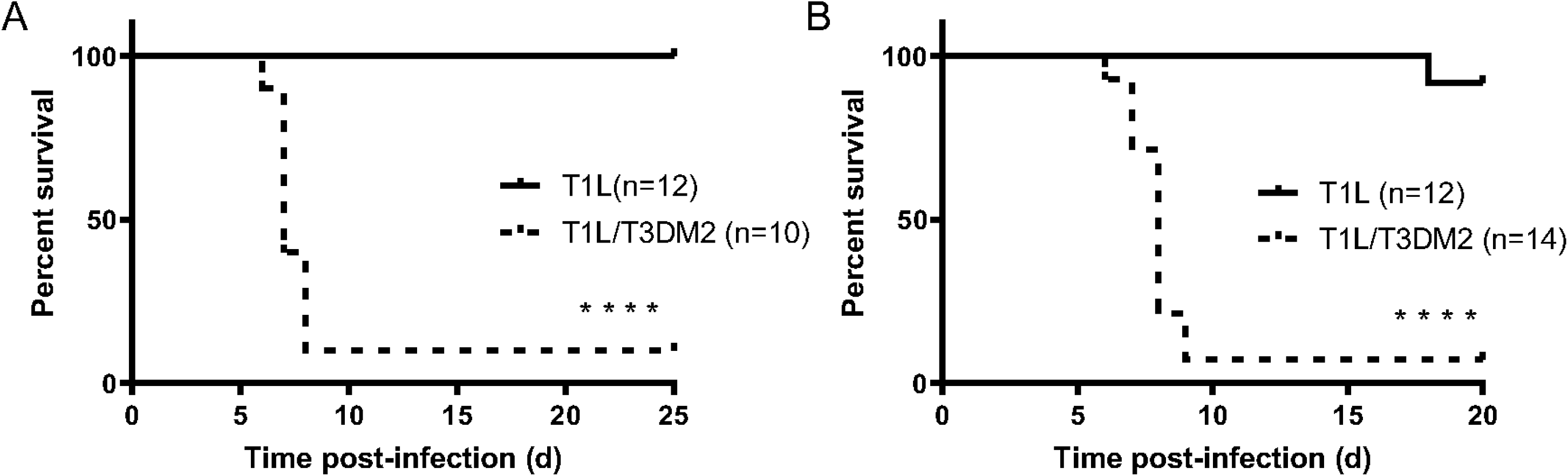
T1L/T3DM2 is more virulent than T1L in neonatal mice. (A) Neonatal C57BL/6 mice were inoculated orally with 10^4^ PFU of T1L or T1L/T3DM2 viruses. Mice were monitored for 25 days. (B) Neonatal C57BL/6 mice were inoculated intracranially with 100 PFU of T1L or T1L/T3DM2 viruses. Mice were monitored for 20 days. ****, *P* < 0.0001 as determined by log-rank test in comparison to T1L.

### T1L/T3DM2 replicates and disseminates more efficiently than T1L *in vivo*

We next measured viral burdens following oral inoculation with T1L or T1L/T3DM2 viruses (Fig 2). At days 1 and 2 post-infection, T1L/T3DM2 produced higher viral titers in the intestine than T1L. However, T1L and T1L/T3DM2 intestinal viral titers equilibrated by day 4. In secondary organs (liver, heart, brain, and spleen), T1L/T3DM2 produced higher viral loads than T1L at most time points and achieved higher peak viral titers than T1L in each organ tested. The largest difference in peak viral titers between T1L/T3DM2 and T1L was observed in the heart, where T1L/T3DM2 viral loads were approximately 50-fold higher than T1L at 8 days post-infection.

**Figure 2.**
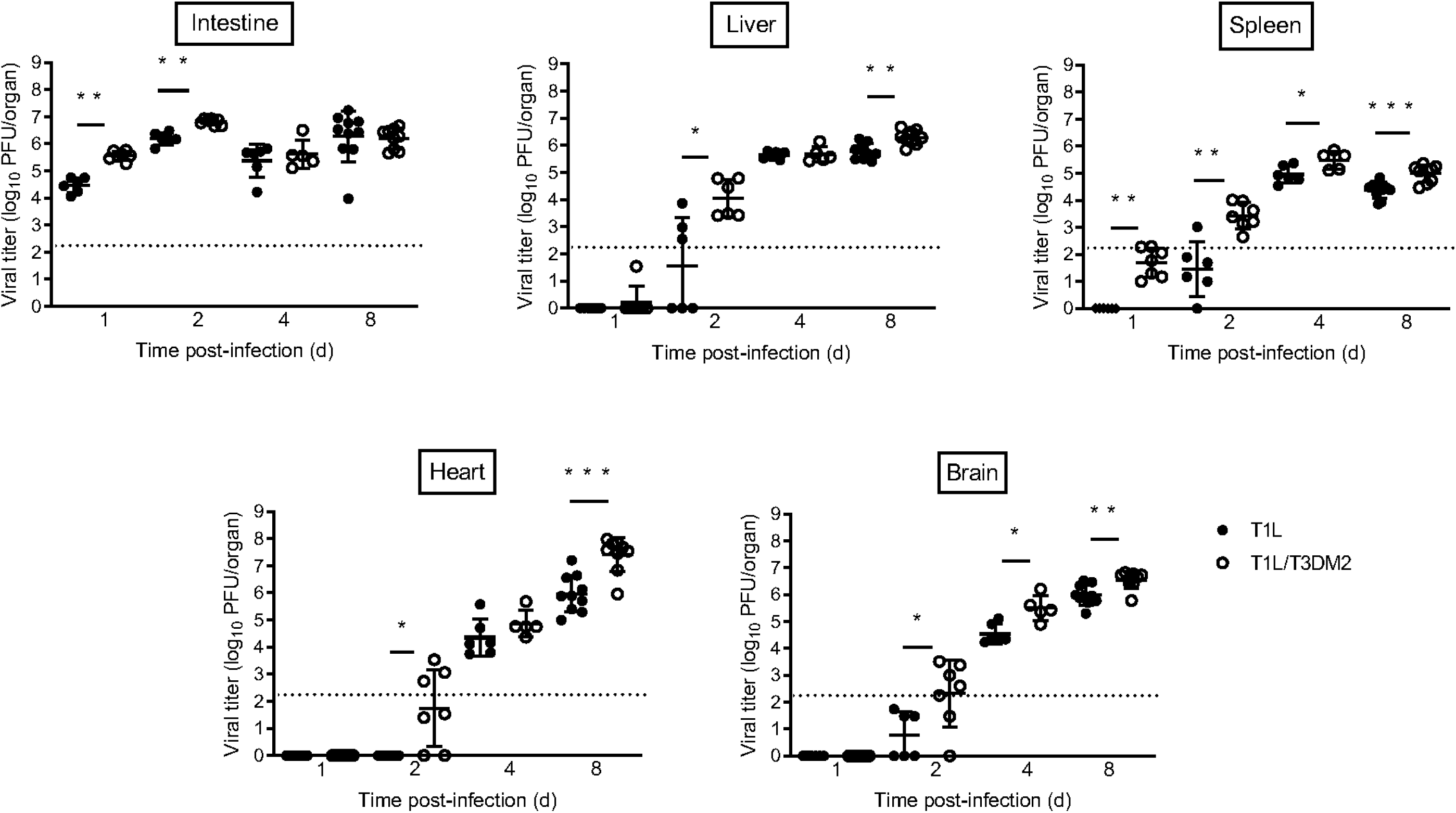
T1L/T3DM2 disseminates more efficiently than T1L following oral inoculation. Neonatal C57BL/6 mice were inoculated orally with 10^4^ PFU of T1L or T1L/T3DM2 viruses. At days 1, 2, 4, and 8, mice were euthanized and the indicated organs were resected. Viral titers in each organ were determined by plaque assay. Error bars represent SD. *, *P* < 0.05; **, *P* < 0.01; ***, *P* < 0.001 as determined by Mann-Whitney test.

We also assessed virulence following intracranial (IC) inoculation. Similar to oral inoculation, T1 reoviruses also disseminate systemically following IC inoculation (28, 29). Like we observed following oral inoculation, T1L/T3DM2 produced higher viral titers at earlier time points and had higher peak viral loads compared to T1L in each organ when virus was introduced IC (Fig 3). Viral titers in the heart were higher following IC inoculation compared to oral inoculation at day 4 (Fig 2 vs Fig 3). Reovirus may disseminate more rapidly after IC inoculation than following oral inoculation, possibly due to the virus needing to overcome fewer physiological barriers when introduced into the brain compared to the intestine. Again, the largest difference in peak viral titers between T1L/T3DM2 and T1L was found in the heart. T1L/T3DM2 viral loads were 10-fold higher than T1L at 8 days post-infection. Together, these data indicate that the M2 allele influences reovirus replication and dissemination *in vivo*.

**Figure 3.**
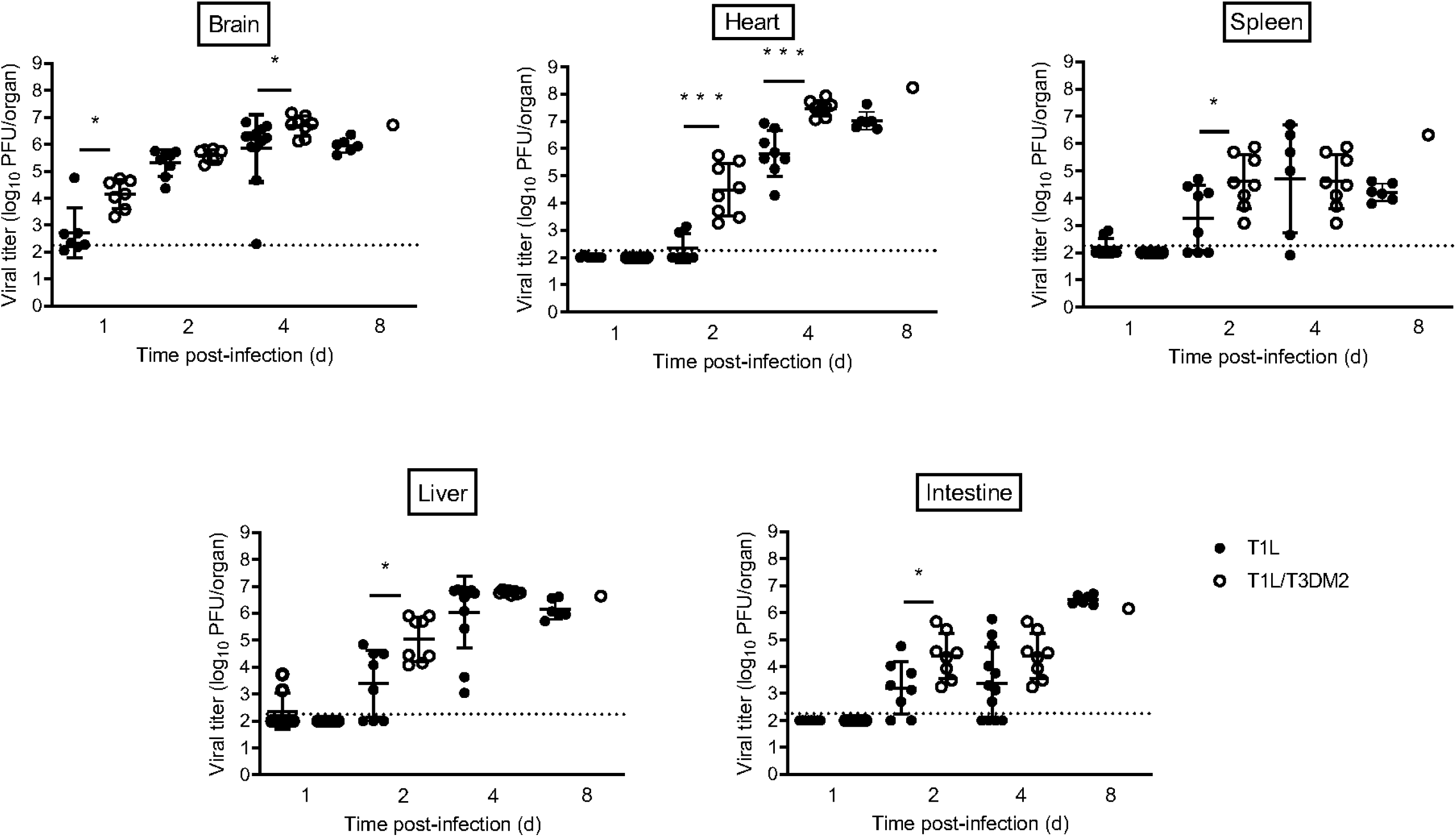
T1L/T3DM2 disseminates more efficiently than T1L following intracranial inoculation. Neonatal C57BL/6 mice were inoculated intracranially with 100 PFU of T1L or T1L/T3DM2 viruses. At days 1, 2, 4, and 8, mice were euthanized and the indicated organs were resected. Viral titers in each organ were determined by plaque assay. Error bars represent SD. *, *P* < 0.05; ***, *P* < 0.001 as determined by Mann-Whitney test.

### Systemic dissemination is required for T3D M2-mediated lethality

All reovirus serotypes require junctional adhesion molecule-A (JAM-A) for systemic dissemination (1, 3, 4). To determine whether systemic spread is required for the virulence differences between T1L and T1L/T3DM2, we measured survival of neonatal JAM-A^−/−^ mice infected with T1L or T1L/T3DM2 viruses (Fig 4). Following oral inoculation, none of the JAM-A^−/−^ mice succumbed to T1L and 10 of 11 (>90%) mice inoculated with T1L/T3DM2 survived (Fig 4A). T1L/T3DM2 produced higher viral titers in the intestine than T1L at days 1 and 2 post-infection, but T1L/T3DM2 and T1L viral loads were equal by day 8 (Fig 4A). Intestinal viral loads in JAM-A^−/−^ mice were comparable to those observed in wild-type mice (Fig 2 vs Fig 4A). This observation is consistent with published results demonstrating that JAM-A is dispensable for reovirus replication in the intestine (30). Low levels of virus were found in the heart and brain from a small number of JAM-A^−/−^ mice but the majority of animals had no detectable virus. Although most JAM-A^−/−^ animals had measurable T1L and T1L/T3DM2 titers in the liver and spleen, viral loads produced in all organ from JAM-A^−/−^ mice were substantially lower than in wild-type mice (Fig 2 vs Fig 4A).

**Figure 4.**
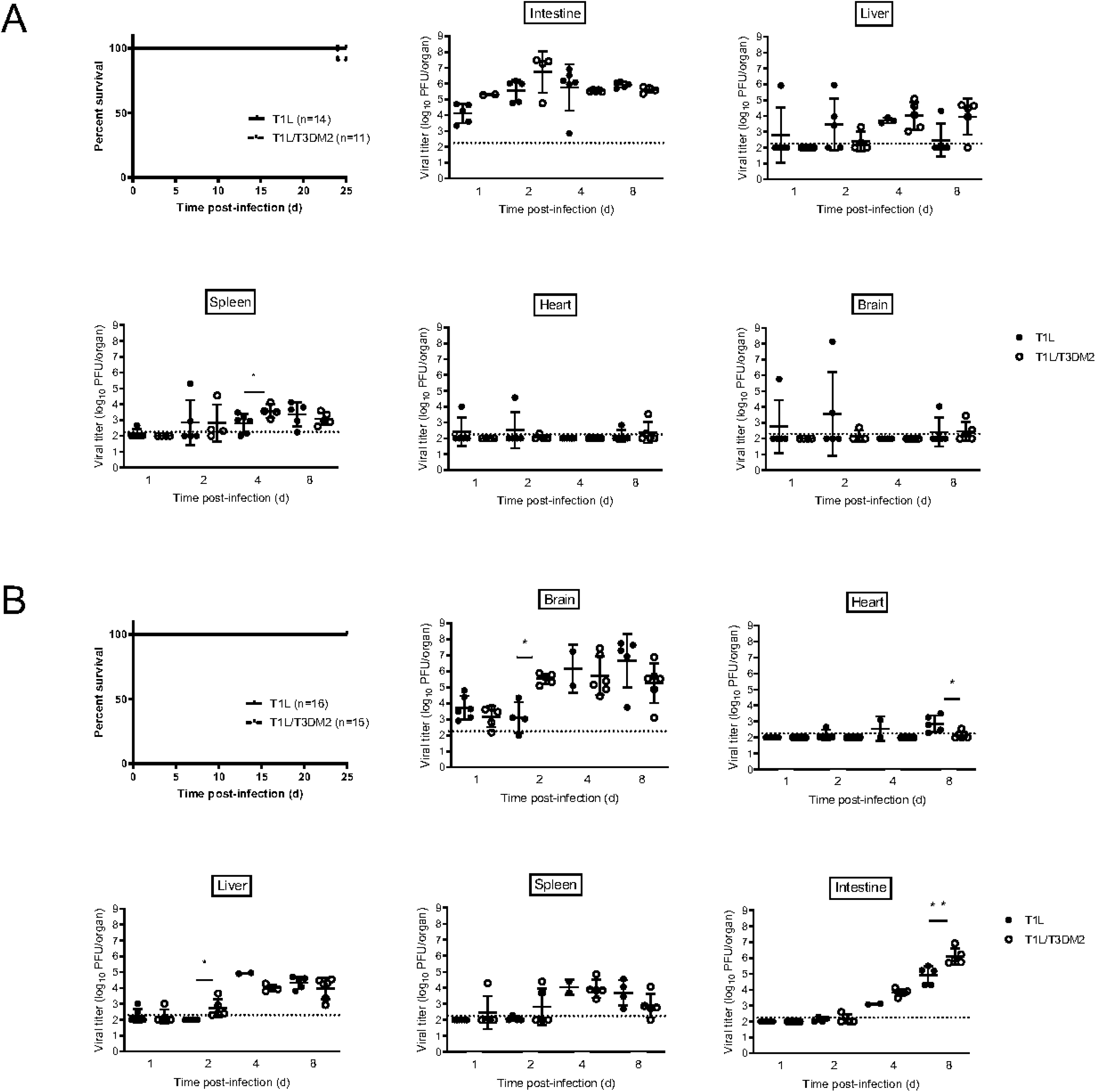
Systemic dissemination is required for T1L/T3DM2 lethality. Neonatal JAM-A^−/−^ mice were inoculated (A) orally with 10^4^ PFU or (B) intracranially with 100 PFU of T1L or T1L/T3DM2 viruses. For survival experiments, mice were monitored for 25 days. For titers experiments, at days 1, 2, 4, and 8, mice were euthanized, and the indicated organs were resected. Viral titers in each organ were determined by plaque assay. Error bars represent SD. *, *P* < 0.05; **, *P* < 0.01 as determined by Mann-Whitney test.

Similar results were obtained following intracranial inoculation. All JAM-A^−/−^ mice infected with T1L or T1L/T3DM2 survived (Fig 4B). These data indicate that JAM-A expression is required for T1L/T3DM2 lethality *in vivo*. T1L and T1L/T3DM2 replicated comparably the brain (Fig 4B), which is consistent with JAM-A being dispensable for reovirus replication in the CNS (40). In secondary organs, T1L and T1L/T3DM2 viral titers were below the level of detection at 1 and 2 days post-infection, but some virus was detected in the heart, liver, and spleen at days 4 and 8 (Fig 4B). However, the viral titers produced in JAM-A^−/−^ mice were substantially lower than the viral loads measured in wild-type animals (Fig 3 vs Fig 4B). T1L and T1L/T3DM2 viral titers in the intestine increased over time, and T1L/T3DM2 produced higher intestinal viral loads than T1L at day 8 (Fig 4B). The dramatic increase in intestinal titers at day 8 may be due to T1L and T1L/T3DM2 establishing viremia in JAM-A^−/−^ mice following IC inoculation, albeit at lower level than in wild-type mice. Once virus reaches the intestine, it can replicate independently of JAM-A to reach peak titers comparable to those observed following oral inoculation (30). Taken together, these data suggest that systemic dissemination is required for the enhanced virulence of T1L/T3DM2 compared to T1L.

### T1L/T3DM2 is more myocarditic than T1L

Regardless of the inoculation route, T1L/T3DM2 was substantially more virulent than T1L in wild-type mice (Fig 1). During necropsy, we observed that hearts from T1L infected mice were largely normal, with a few mice having small cardiac lesions (Fig 5A and B). This result is consistent with published reports that T1L is mildly myocarditic (7, 31, 32). In contrast, hearts from T1L/T3DM2-infected mice were grossly abnormal. In some animals the entire heart appeared opaque, which is indicative of severe myocarditis. These data indicate that substitution of reovirus T3D M2 gene enhances the capacity of T1L to cause cardiac pathology. In JAM-A^−/−^ mice, hearts from T1L or T1L/T3DM-infected animals lacked any signs of inflammation (data not shown). This finding indicates that reovirus dissemination to the heart is required for the development of cardiac lesions upon reovirus infection. It also is possible that JAM-A functions as a reovirus receptor for cardiac cells. Thus, in addition to decreased dissemination in JAM-A-deficient mice, any virus reaching the heart may not be able to infect cardiac cells and cause heart pathology.

**Figure 5.**
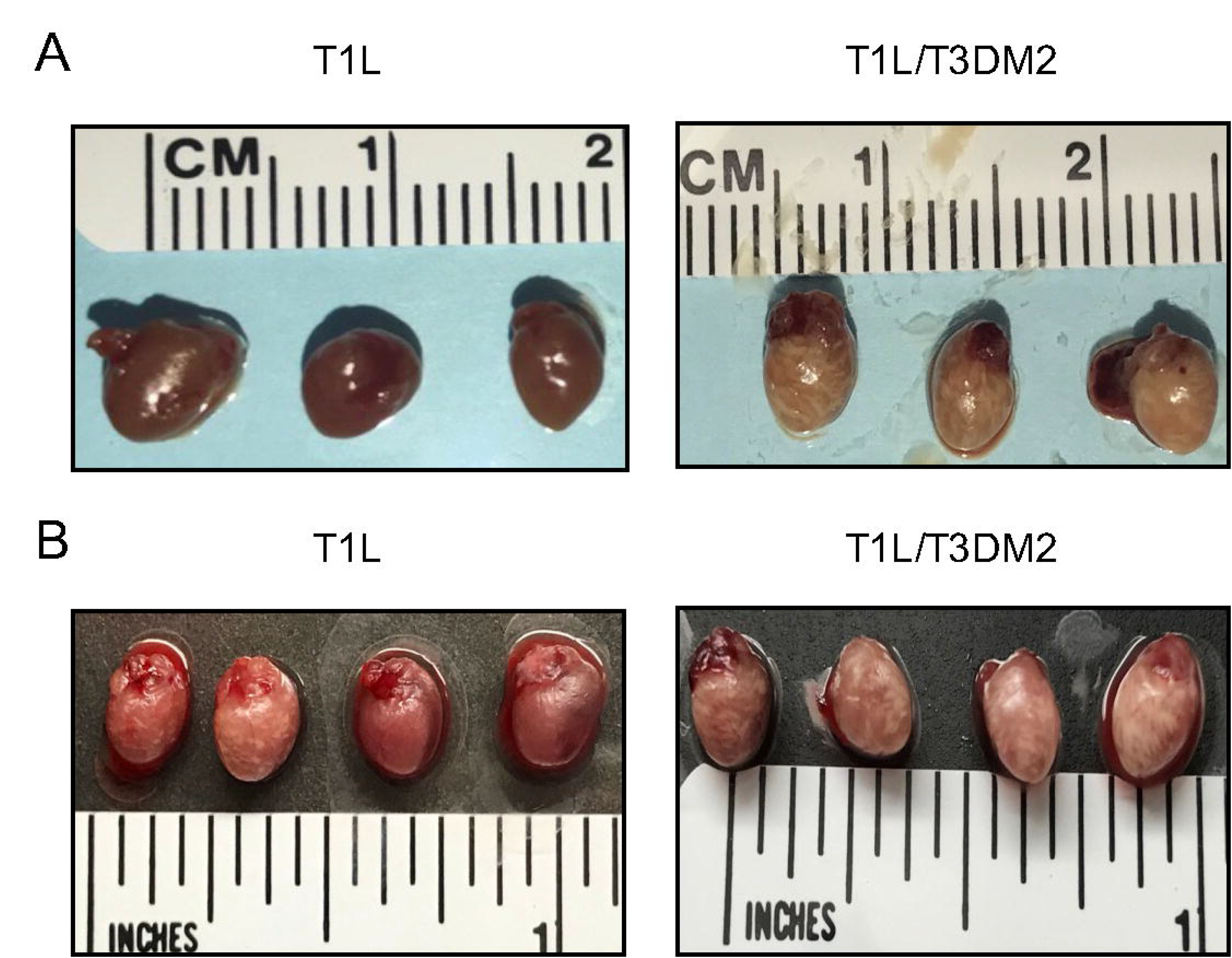
T1L/T3DM2 causes more cardiac pathology than T1L. Neonatal C57BL/6 mice were inoculated (A) orally with 10^4^ PFU or (B) intracranially with 100 PFU of T1L or T1L/T3DM2. Neonatal JAM-A^−/−^ mice were inoculated (C) orally with 10^4^ PFU or (D) intracranially with 100 PFU of T1L or T1L/T3DM2. At 8 days, mice were euthanized and hearts were resected and imaged.

We next performed histological analysis of hearts from wild-type mice infected with T1L or T1L/T3DM2 (Fig 6A). In contrast to mock-infected hearts, T1L-infected hearts had small patches of cardiomyocyte necrosis admixed with pyknotic nuclear debris and histiocytes. T1L lesions typically contained 2-3 cardiomyocytes exhibiting myolysis, which is an indicator of destruction of cardiac muscle tissue. In addition, scant lymphocyte and histiocyte infiltration were observed in the epicardium. These data are consistent with previous observations of T1L inducing mild myocarditis in neonatal wild-type mice. On the other hand, hearts from mice infected with T1L/T3DM2 displayed more frequent cardiomyocyte damage and had increased lymphocytic and histiocytic infiltration in the epicardium compared to T1L-infected hearts (Fig 6A). We quantified the number of inflammatory lesions (Fig 6B), lesion size (Fig 6C), and total percentage of the heart with infiltrating immune cells (Fig 6D) in H&E-stained sections hearts from mock-, T1L-, and T1L/T3DM2-infected mice. There was no difference in the number of cardiac lesions between T1L- and T1L/T3D/M2-infected hearts. This finding suggests that T1L and T1L/T3DM2 initiate infection of the heart at a similar rate. Cardiac lesions from mice infected with T1L/T3DM2 were larger, and the overall level of cardiac immune cell infiltration was significantly higher, than in T1L-infected mice (Fig 6C). Hearts from mice infected with T1L/T3DM2 showed markedly more viral antigen staining compared to T1L (Fig 6B). This observation is consistent with the higher peak titers produced by T1L/T3DM2 compared to T1L in the heart at day 8 (Fig 2 and Fig 3). It is of note that regions of viral antigen staining in T1L and T1L/T3DM2-infected hearts overlapped with myocardial lesions (Fig 6B). This finding indicates that reovirus replication in cardiomyocytes is associated with lesion formation (11). Lesions from T1L- and T1L/T3DM2-infected hearts also contained cells that stained positive for activated caspase-3, which is a marker of apoptosis (Fig 6C). Together, these data indicate that T1L- and T1L/T3DM2-induced cardiac lesions contain reovirus-infected cells, infiltrating immune cells, and apoptotic cells, with T1L/T3DM2 causing more extensive lesion formation than T1L.

**Figure 6.**
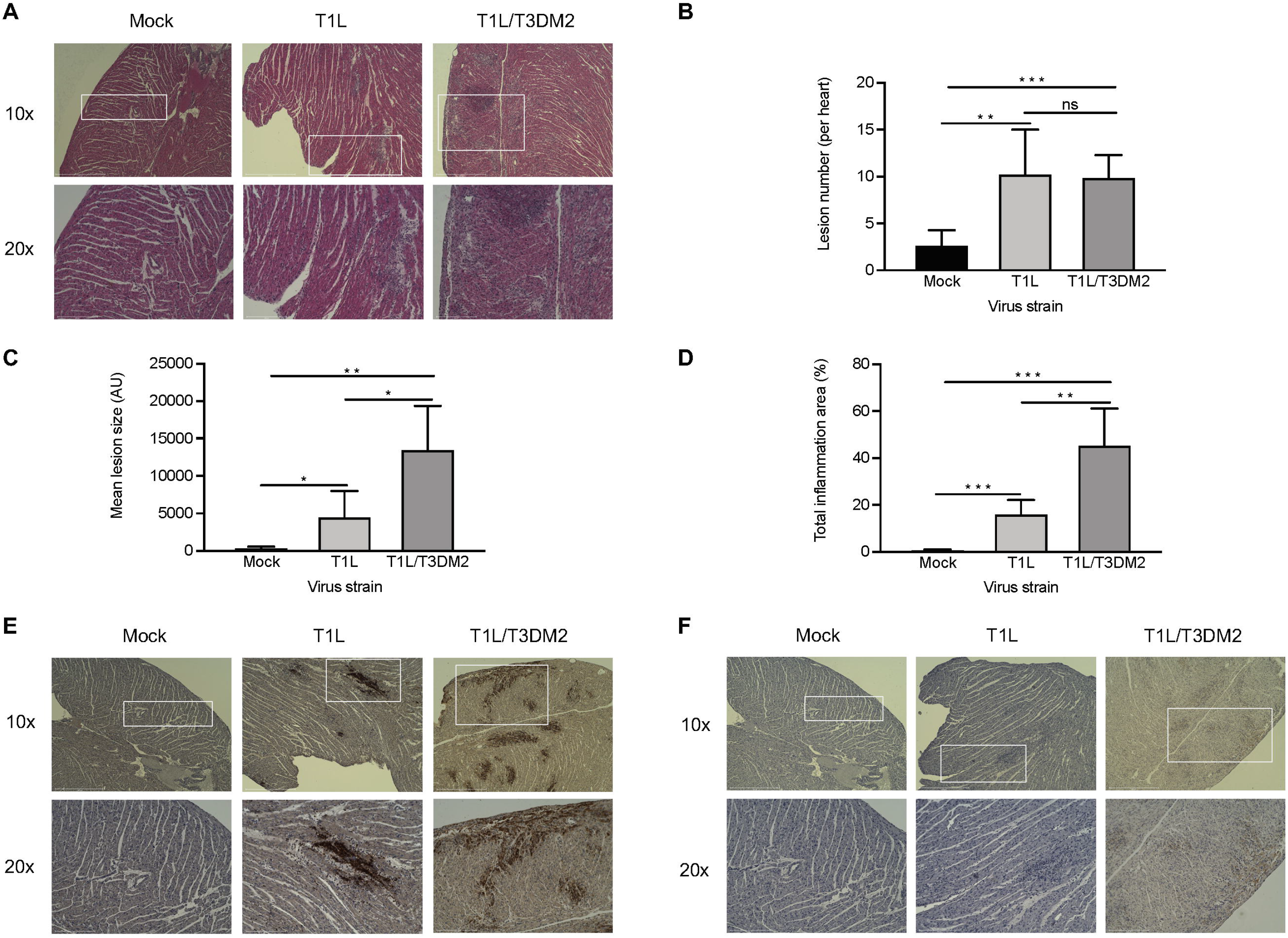
T1L/T3DM2 is more myocarditic than T1L. Neonatal C57BL/6 mice were mock-infected with PBS or inoculated orally with 10^4^ PFU of T1L or T1L/T3DM2 viruses. At 8 days, the mice were euthanized, the hearts were resected, paraffin embedded, and sectioned. Consecutive sections were stained with (A) hematoxylin and eosin (H&E), (E) polyclonal reovirus antiserum, or (F) cleaved-caspase-3 antibodies. Sections were imaged at 10x magnification (top panels) or 20x (bottom panels). Using H&E stained sections, (B) cardiac lesion number, (C) lesion size (in arbitrary units; AU), and (D) total area of inflammation (expressed as percentage of the total heart) were quantified (*n* = 5 mice per group). Error bars indicate SD. *, *P* < 0.05; **, *P* < 0.01 ***, *P* < 0.001 (as determined by Student’s *t*-test).

## DISCUSSION

In this study, we found that the M2 gene is a determinant of reovirus myocarditis. Reovirus strain T1L is non-lethal in neonatal mice although it induces mild heart inflammation in a subset of animals. However, replacement of the T1L M2 gene with the T3D M2 allele (T1L/T3DM2) was sufficient to confer a lethal phenotype that is characterized by marked inflammation in nearly 100% of infected animals. T1L/T3DM2 replicated to significantly higher titers at initial site of inoculation (Fig 2 and Fig 3), disseminated to secondary organs with more rapid kinetics than, and produced higher peak organ titers than T1L (Fig 2 and Fig 3). T1L/T3DM2 binds and infects L929 cells more efficiently than T1L, leading to a higher percentage of cells becoming infected and ultimately production of higher progeny yields than T1L (20). It is possible T1L/T3DM2 initially infects more cells than T1L, leading to higher progeny yields at the primary site of replication that disseminate systemically and infect target organs more efficiently, including the heart. The greatest difference in peak viral titers between T1L and T1L/T3DM2 was observed in the heart, where T1L/T3DM2 viral progeny were approximately 50-fold and 10-fold higher than T1L following oral and intracranial inoculation, respectively (Fig 2, Fig 3). Previous studies indicate that reovirus replication levels in the heart do not correlate with myocarditis induction (10). It is possible that delivery of T1L/T3DM2 virus to the heart more rapidly than T1L allows T1L/T3DM2 to reach a threshold that contributes to the myocarditic capacity of T1L/T3DM2.

Apoptosis is a key pathogenic mechanism that contributes to reovirus myocarditis (6, 33). Clone 8B induces caspase-3 activation within the myocardium of infected hearts, enhancing myocardial injury (34). Moreover, myocarditis development is dramatically reduced in caspase-3-deficient mice and mice treated with caspase inhibitors. (34–36). Reoviruses display serotype-specific differences in apoptosis induction, with T3 reoviruses inducing substantially more apoptosis than T1 strains (18). A determinant of reovirus apoptosis is the M2 gene (6, 18, 27). We found that hearts from mice infected with T1L/T3DM2 had more prominent activated caspase-3 staining than T1L-infected hearts (Fig 6). Therefore, we hypothesize that the T3D M2 allele enhances apoptosis in infected cells, which promotes myocardial injury and inflammation, which is the hallmark of myocarditis.

The inflammatory response also may contribute to the enhanced myocarditic capacity of the T1L/T3DM2 virus. The adaptive immune response is dispensable for reovirus-induced myocarditis (6, 37). However, innate immune cells may contribute to the heart inflammation and cell death observed following infection with the T1L/T3DM2 virus. Similar to what we observed for T1L/T3DM2, clone 8B induced significant mononuclear cardiac infiltration (11). The myocarditic reovirus strain T1L/T3Dμ1σ3, which contains the M2 and S4 genes from T3D, elicits substantial pro-inflammatory cytokine production in the heart that promotes cardiomyocyte damage (12).

T1L is mildly myocarditic, whereas T3D is non-myocarditic (6, 7, 31). However, reassortant viruses generated from cells coinfected with T1L and T3D have the capacity to cause severe myocarditis (11, 38). A number of highly myocarditic reassortants contain the T3D M2 gene. Clone 8B contains the S1 and M2 genes from T3D, while clone KC19 contains the T3D S1, M2, and S3 genes (7, 11). Clone EW60 is genetically similar to the T1L/T3DM2 used in this study as it is a single-gene reassortant containing the T3D M2 gene in a T1L genetic background (31). Clone E3, a single gene reassortant with the T1LM2 gene in an otherwise T3D genetic background, is non-myocarditic in neonatal mice (8). Together, these findings suggest that reovirus myocarditic capacity can be modulated by the M2 gene. Here, we used T1L/T3DM2 produced using reverse genetics to clarify and characterize how M2 contributes to reovirus myocarditis. However, the M2 gene is not the sole determinant of reovirus myocarditis. The M1 gene, which encodes inner core protein μ2 that functions as a co-factor for the reovirus RNA-dependent RNA polymerase, also is a determinant of reovirus myocarditis (6, 8, 10, 32). M1 enhances reovirus myocarditis by potentiating replication in cardiac cells and modulating the sensitivity to type-1 interferon (8, 39). In addition, distinct T3D clones have evolved as the virus is passaged by different laboratories and these clones can differ phenotypically (40–42). Comparing the myocarditic capacity of M2 genes from different T3D clones may provide insight into how M2 causes myocarditis. There may not be a clear-cut mechanism underlying how T1L/T3DM2 enhances myocarditis. It is possible that enhanced dissemination to, and replication in, the heart coupled with amplified apoptotic capacity combine to increase the level of cardiac damage induced by T1L/T3DM2. In turn, the recruitment of immune cells to fight the infection may further exacerbate cardiac damage by production of pro-inflammatory cytokines and directly or indirectly killing cells.

Other models of virus-induced myocarditis share similarities with reovirus (43–46). Severe acute respiratory syndrome coronavirus-2 (SARS-CoV-2) replication in the myocardium is hypothesized to drive the pathological changes observed in the heart tissue (47). Coxsackievirus B3 and murine cytomegalovirus (CMV) myocarditis require viral replication and apoptosis induction (43–46). Studies of how reovirus causes pathology in cardiac muscle cells can provide valuable information about how the heart combats viral infection.

## MATERIALS AND METHODS

### Cells

Spinner-adapted murine L929 fibroblasts were maintained in Joklik’s modified minimum essential medium (JMEM, Sigma) supplemented with 5% heat-inactivated fetal bovine serum (FBS, Invitrogen), 1% 200 mM L-glutamine (Invitrogen), 1% 10,000 U/ml penicillin/ 10,000 μg/ml streptomycin (Invitrogen), and 0.1% 250 μg/ml amphotericin B (Sigma).

### Viruses

Recombinant reoviruses (T1L and T1L/T3DM2) were generated using plasmid-based reverse genetics (48, 49). Reovirus virions were purified from second- or third-passage L929 cell lysates infected with twice-plaque-purified reovirus (50). Reovirus particles were Vertrel (TMC industries) extracted from infected cell lysates, layered onto 1.2-to 1.4-g/cm^3^ CsCl gradients, and centrifuged at 107,240 × *g* for 18h or 144,302 × *g* for 5h. Virions were collected and dialyzed against dialysis buffer (150 mM NaCl, 15 mM MgCl, and 10 mM Tris-HCl [pH 7.4]). Viral titers were determined by plaque assay using L929 fibroblasts (51). To confirm the viral gene segments, viral dsRNA was purified from virions and separated by SDS-PAGE. Gene segments were visualized by ethidium bromide staining (48).

### Mouse experiments

C57BL/6 mice were obtained from Jackson Laboratory. C57BL/6-JAM-A^−/−^ mice were obtained from the Dermody Laboratory at the university of Pittsburgh. Animal husbandry and housing were performed following the guidelines of the Division of Laboratory Animal Medicine (DLAM) at the University of Arkansas for Medical Sciences (UAMS). Neonatal mice (3-5 days old) were infected orally with 10^4^ PFU or intracranially with 100 PFU of rsT1L or rsT1L/T3DM2. For survival experiments, infected mice were monitored for 20 or 25 days and sacrificed if moribund. Brain, heart, spleen, liver, and intestine organs were resected at the times indicated in the figures legends, homogenized, and viral titers were determined by plaque assay on L929 cells. Resected hearts were imaged at the times indicated in the figure legends using a hand-held camera.

### Histology and immunohistochemistry

Neonatal C57BL/6 mice (3-5 days old) were mock-infected (PBS) or infected orally with 10^4^ PFU of rsT1L or rsT1L/T3DM2. At 8 days post-infection, mice were euthanized and the hearts were resected. Vertical longitudinal cuts of the infected hearts were prepared and fixed in methanol Carnoy’s for 24h followed by 70% ethanol until embedding. The fixed hearts were submitted to the UAMS Experimental Pathology Core Laboratory for paraffin embedding and sectioning (4 μm thickness). Cardiac sections were stained with hematoxylin and eosin (H&E) and a pathologist blinded to the identity of the inoculum examined them under a light microscope. ImageJ software was used to count the number of cardiac lesion as defined as distinct area of immune infiltration. For each cardiac section (10×magnification), the total heart surface area was determined using the HSB threshold color tool and measured in arbitrary units (pixels). Using the Magic Wand tool in ImageJ, each area of immune cells aggregate was selected as a cardiac lesion and the size was measured in arbitrary unit (pixels). The total area of inflammation was calculated as a percentage of the total heart section. Results were collected from 5 hearts per infection. Consecutive sections were processed for immunohistochemical staining with reovirus-specific rabbit polyclonal antiserum (1:1000) or activated caspase-3 (Abcam, 1:100). Images were taken on an EVOS FL Auto 2 (Invitrogen) using the 10x and 20x objectives (133x and 266x total magnification, respectively).

### Statistics

Statistical analysis was performed using Prism software (GraphPad Software Inc.). Differences in survival were determined by Log-rank test. Differences in mean viral titers were determined using Mann-Whitney non-parametric *t*-test. Differences in cardiac lesion number, size, and total area of inflammation were evaluated using an unpaired student *t*-test. *P* values less than 0.05 were considerate statistically significant.

## ACKNOWLEDGMENTS

We thank the Dermody Lab (University of Pittsburgh) for providing the JAM-A^−/−^ mice. We are grateful to Jennifer James at the UAMS Experimental Pathology Core (UAMS) for assistance with histological sample preparations. This research was supported by Public Health Award R01 AI118801 (K.W.B.) and R01 AI110637 (P.D). Additional support was provided by the Center for Microbial Pathogenesis and Host Inflammatory Response (P20 GM103625).

